# Is mRNA decapping activity of ApaH like phosphatases (ALPH’s) the reason for the loss of cytoplasmic ALPH’s in all eukaryotes but Kinetoplastida?

**DOI:** 10.1101/2020.12.17.423368

**Authors:** Paula Andrea Castañeda Londoño, Nicole Banholzer, Bridget Bannermann, Susanne Kramer

## Abstract

**Background:** ApaH like phosphatases (ALPHs) originate from the bacterial ApaH protein and are present in eukaryotes of all eukaryotic super-groups; still, only two proteins have been functionally characterised. One is ALPH1 from the Kinetoplastid *Trypanosoma brucei* that we recently found to be the mRNA decapping enzyme of the parasite. mRNA decapping by ALPHs is unprecedented in eukaryotes, which usually use nudix hydrolases, but the bacterial ancestor protein ApaH was recently found to decap non-conventional caps of bacterial mRNAs. These findings prompted us to explore whether mRNA decapping by ALPHs is restricted to Kinetoplastida or more widespread among eukaryotes.

**Results:** We screened 824 eukaryotic proteomes with a newly developed Python-based algorithm for the presence of ALPHs and used the data to refine phylogenetic distribution, conserved features, additional domains and predicted intracellular localisation of ALPHs. We found that most eukaryotes have either no ALPH (500/824) or very short ALPHs, consisting almost exclusively of the catalytic domain. These ALPHs had mostly predicted non-cytoplasmic localisations, often supported by the presence of transmembrane helices and signal peptides and in two cases (one in this study) by experimental data. The only exceptions were ALPH1 homologues from Kinetoplastida, that all have unique C-terminal and mostly unique N-terminal extension, and at least the *T. brucei* enzyme localises to the cytoplasm. Surprisingly, despite of these non-cytoplasmic localisations, ALPHs from all eukaryotic super-groups had *in vitro* mRNA decapping activity.

**Conclusions:** ALPH was present in the last common ancestor of eukaryotes, but most eukaryotes have either lost the enzyme since, or use it exclusively outside the cytoplasm in organelles in a version consisting of the catalytic domain only. While our data provide no evidence for the presence of further mRNA decapping enzymes among eukaryotic ALPHs, the broad substrate range of ALPHs that includes mRNA caps provides an explanation for the selection against the presence of a cytoplasmic ALPH protein as a mean to protect mRNAs from unregulated degradation. Kinetoplastida succeeded to exploit ALPH as their mRNA decapping enzyme, likely using the Kinetoplastida-unique N- and C-terminal extensions for regulation.

## BACKGROUND

Eukaryotic phosphatases play essential roles in regulating many cellular processes and can be classified in several ways, based on catalytic mechanism, substrate specificity, ion requirements and structure. One four-group classification distinguishes phosphoprotein phosphatases (PPPs), metal-dependent protein phosphatases (PPM, sometimes classified as a subgroup of PPP), protein tyrosine phosphatases and aspartic acid-based phosphatases [1]. The eukaryotic PPP group includes the Ser/Thr phosphatases PP1, PP2A, PP2B, PP4, PP5, PP6 and PP7 [1,2] but has been extended to include three families of bacterial origin: Shewanella-like SLP phosphatases, Rhizobiales-like (RLPH) phosphatases and ApaH-like (ALPH) phosphatases [3–6].

ApaH like phosphatases (ALPHs) evolved from the bacterial ApaH protein and were present in the last common ancestor of eukaryotes [3,5]. They are widespread throughout the entire eukaryotic kingdom, albeit have been lost in certain sub-branches such as land plants and chordates [6]. To the best of our knowledge, only two ALPHs have been functionally characterised to date. One is the *S. cerevisiae* ALPH protein (YNL217W), an Zn^2+^ dependent endopolyphosphatase of the vacuolar lumen [7] that is also active with Co^2+^ and possibly Mg^2+^ [8]. The enzyme’s main function is the cleavage of vacuolar poly(P). A function of Ppn2 in stress response is suggested by the fact that the increase in short polyphosphate upon Ppn2 overexpression correlates with an increased resistance to peroxide and alkali [9]. The second characterised ALPH protein is ALPH1 of the Kinetoplastida *Trypanosoma brucei*: we recently found that ALPH1 is the only or major trypanosome mRNA decapping enzyme [10], the enzyme that removes the m^7^ methylguanosine (m^7^G) cap present at the 5’end of most eukaryotic mRNAs. This finding was surprising, as all other known mRNA decapping enzymes belong to a different enzyme family, the nudix hydrolases (the prototype is Dcp2). Trypanosomes lack orthologues to Dcp2 and all decapping enhancers, the likely reason why they use the ApaH like phosphatase ALPH1 instead.

Interestingly, recent data indicate, that the bacterial precursor protein of ALPH, ApaH, may have an analogous function to Trypanosome ALPH1 in decapping of bacterial mRNAs. *In vitro,* ApaH cleaves diadenosine tetraphosphate (Ap4A) into two molecules of ADP [11,12], but is also active towards other NpnN nucleotides (with n?3) [13–15]. Deletion of the ApaH gene *in vivo* causes a marked increase in Np4A levels and a wide range of phenotypes [16–23]. Np4A was therefore suggested to act as an alarmone (a signalling molecule involved in stress response), but no Np4A receptor has yet been identified. Instead, recent data show that stress induced increase in Np4A levels cause massive nucleoside-tetraphosphate capping of bacterial mRNAs [24] mostly or entirely caused by usage of Np4A as non-canonical transcription initiation nucleotide [25]. Many other dinucleoside polyphosphates can be used for co-transcriptional capping even in the absence of stress, including methylated versions [26]. ApaH is the major decapping enzyme for all dinucleoside polyphosphate caps [24,26], suggesting that its main function is the regulation of gene expression via regulating mRNA decapping. Intriguingly, the enzyme can both enhance decapping (by decapping nucleoside-tetraphosphate capped RNA) and inhibit decapping (by cleaving Np4A and preventing its incorporation to the mRNA).

Puzzled by these novel functions of both bacterial ApaH and trypanosome ALPH1 in mRNA decapping, we here set out to investigate, whether mRNA decapping is a major function of eukaryotic ALPHs, or whether this function is restricted to trypanosomes. We developed a Python algorithm for the identification of ALPHs in all available eukaryotic reference proteomes and identified 412 ALPHs in 324/824 proteomes. To our surprise, almost all ALPH proteins consisted exclusively of the catalytic domain and almost all had predicted transmembrane regions or signal peptides and predicted non-cytoplasmic localisation. Among the few exceptions were all orthologues to trypanosome ALPH1 present in all organisms of the Kinetoplastida group: these had C-terminal and, with only 2 exceptions N-terminal extensions and cytoplasmic localisation. The absence of any (regulatory) domains and the non-cytoplasmic localisation predictions argue against mRNA decapping being a widespread function of eukaryotic ALPHs outside the Kinetoplastida. We tested three randomly chosen ALPH proteins of three different eukaryotic super-groups for mRNA decapping activity *in vitro*, and all were active, indicating a broad substrate range of this enzyme class. The mRNA decapping activity of ALPHs provides a possible explanation for the evolutionary selection against the presence of a cytoplasmic ALPH in eukaryotes by either losing the gene, or targeting the protein to organelles, as an effective mean to protect mRNAs from unregulated degradation. Only trypanosomes have succeeded to exploit the mRNA decapping activity of ALPH to their advantage, by adding regulatory domains, likely involved in regulating enzyme activity.

## RESULTS

### Identification of ApaH like phosphatases in available eukaryotic proteomes

ALPHs belong to the PPP family of phosphatases and possess the four conserved signature motifs (motif 1-4) of this family, GDxHG, GDxxDRG, GNHE, and HGG, sometimes with conservative substitutions [2]. One distinctive feature of ALPHs are two changes in the GDxxDRG motif: The second Asp is replaced by a neutral amino acid and the Arg residue is replaced by Lys. In addition, ALPHs have two C-terminal motifs [3,6] that we here call motif 5 and 6. We screened 824 complete eukaryotic proteomes (Table S1a) for the presence of ApaH like phosphatases with a home-made Python algorithm; these included all reference proteomes available on UniProt [27] and all available Kinetoplastida proteomes available on TriTrypDB [28,29]. The algorithm is based on recognising the six sequence-motifs characteristic for ALPHs. The matrices used to define these motifs were stepwise optimised on yeast and Kinetoplastida proteomes to not miss any ALPH (controlled by BLAST) while on the other hand not to recognise PPPs, and in particular not the closely related phosphatases SLP, RLPH and ApaH (sequences taken from [6]). The final algorithm also included restrictions on distances between the motifs 1 and 2, 2 and 3 and 5 and 6 that we found to be highly conserved. BLAST screens on selected proteomes without ALPH revealed that ALPHs were either fully absent or present in a truncated version missing at least one of the motifs, often the most N or C-terminal one. Such truncated ALPHs were not included to the final list, even though a subset may have arisen from sequencing or annotation errors. Only for the Kinetoplastida, ALPHs with wrongly annotated start codons were manually included (based on comparison with related Kinetoplastida).

Figure 1 summarises all organisms of this study in phylogenetic groups based on the latest eukaryotic classification suggested by [30]. For each group, the fraction of organisms with and without ALPH is indicated in orange and blue, respectively. 324 of all 824 organisms included in this study have at least one ALPH, and these organisms are distributed in a patchy way throughout all eukaryotes. Most Euglenozoa have ALPHs, many Stramenopiles and Dikarya, but also Rhodophyceae and Chloroplastida and many Metazoa. ALPHs are absent from land plants, Apicomplexa and Ciliata. They are largely absent from Chordata (with the exception of *Branchiostoma floridae)* and from Ecdysozoa (with 4 exceptions) and fully absent from the few available proteomes of Amoebozoa and Metamonada. A phylogenetic tree built from the catalytic domains of ALPHs mostly reflects the eukaryotic tree (Supplementary Figure S1 and Supplementary Table S1b). Taken together, the data indicate that ALPH was present in the last common ancestor of all eukaryotes and was then selectively lost in certain sub-branches. Our data extend and agree to the data of [6].

**Figure 1:**
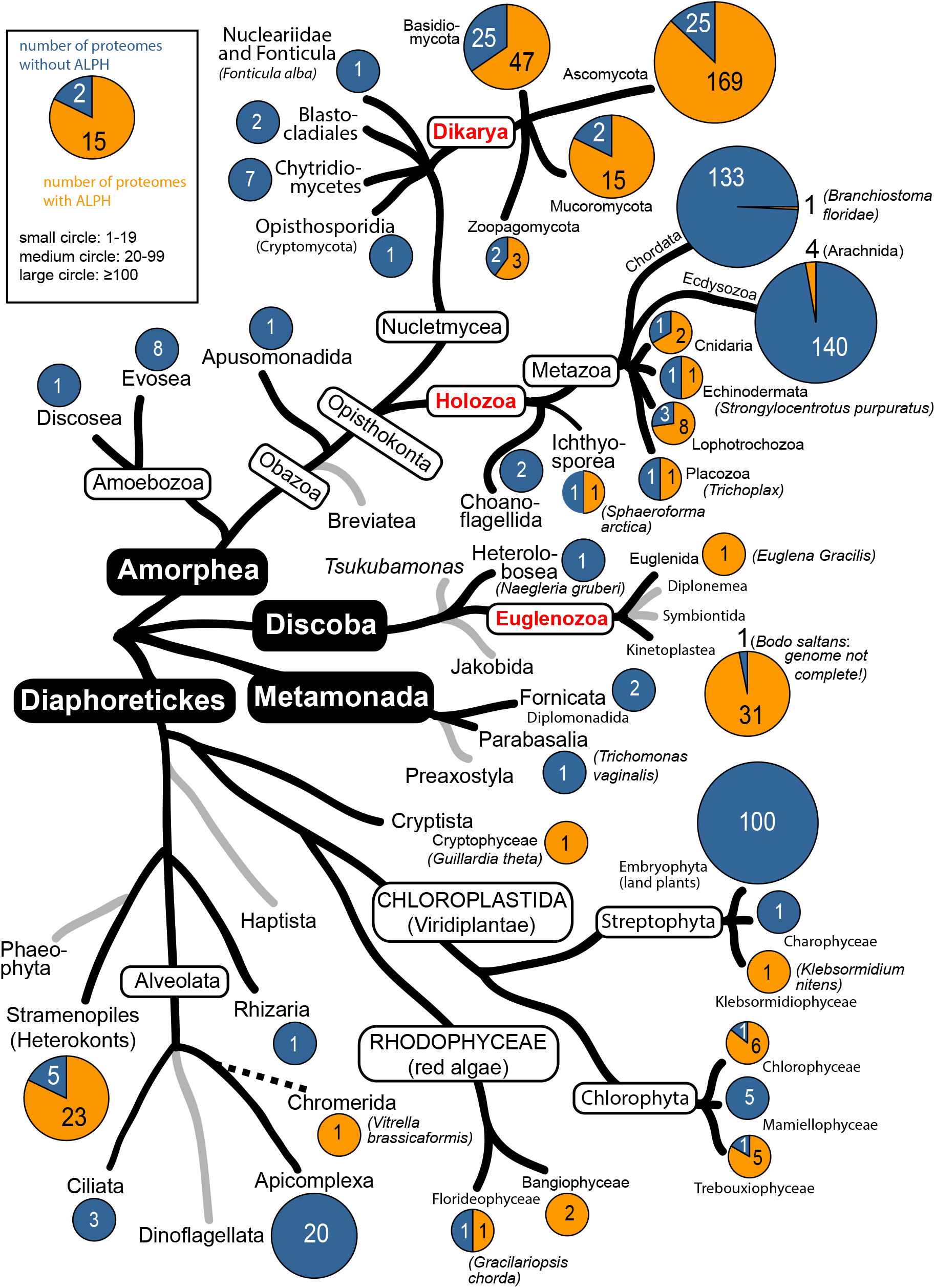
Presence and absence of ALPH in the different eukaryotic subgroups. 824 eukaryotic proteomes were screened for the presence of ALPH. Absence or presence of ALPH is shown in pie diagrams in blue and orange, respectively, for each phylogenetic group as indicated. The diameter of a circle roughly reflects the number of available proteomes. Information for the phylogenetic tree was taken from [30]. All organisms can be found in Supplementary Table S1.

The aim of this work was not to analyse horizontal gene-transfer between eukaryotes, bacteria and archaea; however, we could confirm the presence of ALPHs in a subgroup (11/25) of *Myxococcales,* as described in [6] (Supplementary Table S2A) and we detected ALPH in 1/285 archaean proteomes (OX=1906665 GN=EON65_52185, UniProt: UP000292173) (Supplementary Table S2B). All prokaryotic ALPHs consisted mostly of the catalytic domain with almost no N- or C terminal extensions.

### General features of ApaH-like phosphatases

The dataset was used to refine the characteristics of ALPH proteins (Figure 2). 39% of all analysed organisms have at least one ALPH isoform. The highest percentage is found in Discoba with 94%, the lowest in Diaphoretickes with 22% (Figure 2A). Of the 324 APLH-positive organisms, 20% have more than one ALPH isoform: most (77%) have two and with one exception no organism has more than four (Figure 2B). Organisms with multiple ALPHs were with 47% mostly enriched among the Discoba (Figure 2B). The vast majority of all ALPH proteins is very short and consist mostly of the catalytic domain (Figure 2C). The median size of the C-termini is with 26 amino acids very short. Only 49 ALPHs have C-termini longer than 100 amino acids and most of these (31) are ALPHs of Discoba. The size of the N-terminus is slightly more variable and has a median of 88 amino acids. Only 58 N-termini are longer than 200 amino acids and many of these (28) belong to ALPHs of Discoba. The largest variance in the size of ALPH N- and C-termini is found in Discoba, reflecting the presence of two very different ALPH variants in the Kinetoplastida (discussed below).

**Figure 2:**
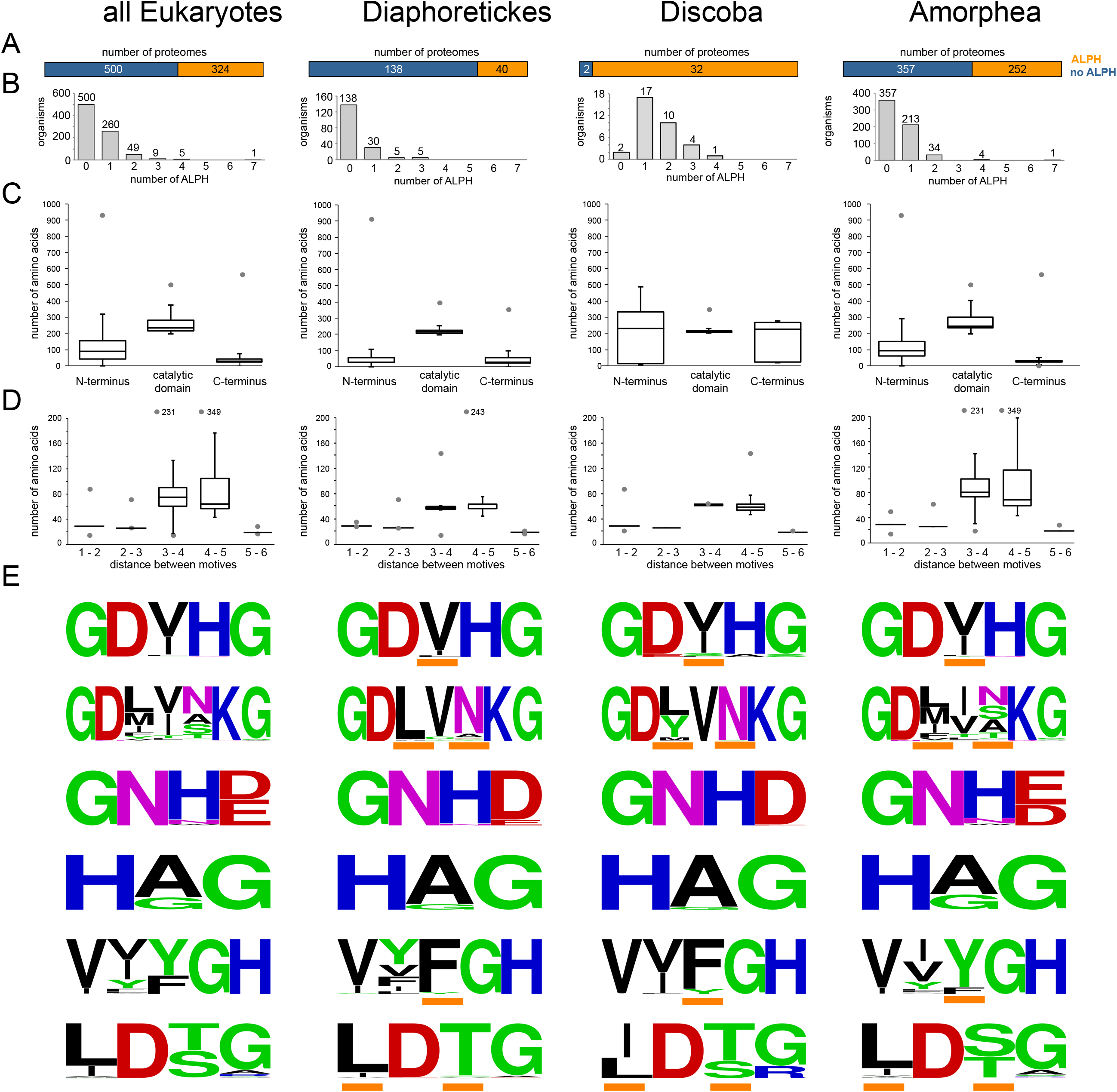
General Features of ALPHs. **(A)** Proteomes with (orange) and without (blue) ALPHs. **(B)** Number of ALPH proteins per organism. **(C)** Sizes of the different ALPH ‘domains’ (N-terminus, catalytic domain, C-terminus) are presented as box plot (waist is median; box is IQR; whiskers are ±1.5 IQR; only the smallest and largest outliers are shown). The catalytic domain is defined as the range between the first and last motif, with an additional six N-terminal amino acids and an additional eight C-terminal amino acids. **(D)** Distances between the six different motifs (in amino acids) are presented as box plot. **(E)** Sequence motifs were created with WebLogo [31]. The most obvious differences between the three groups are marked with orange bars.

The amino acid distances between some of the ALPH motifs were highly conserved (Figure 2D): 95.2 of all ALPHs have between 28 and 30 amino acids between motif 1 and 2. The distance between motif 2 and 3 is 26 amino acids for 83.1% of ALPHs and between 25 and 33 for 98.1% of ALPHs. The distance between the two C-terminal motifs 5 and 6 is 19 amino acids for 93.2% of ALPHs. The distances between motif 3 and 4 and between motif 5 and 6 are less well conserved within all eukaryotes or within Amorphea, but well conserved within the groups of Diaphoretickes and Discoba. Sequence motifs were created for all six motifs (Figure 2E) [31]. Mostly, these are conserved with some group-specific preferences at certain positions indicated with orange bars (Figure 2E).

### ALPHs of Opisthokonts (Dikarya and Holozoa)

The majority of available ALPH sequences (286) are from Dikarya, because the number of available proteomes is high (288) and 81% of these proteomes contain at least one ALPH. In this paper, we include Mucoromycota and Zoopagomycota to the Dikarya as suggested by [30] and indicated in Figure 1. We investigated these ALPH sequences further by looking for predicted domains (Interpro [32]), signal peptides and trans-membrane helices (Phobius [33]) and predicted localisation (WoLF PSORT [34]) (Fig. 3A and Table S1c). ALPHs of Dikarya have very short C-termini (median=26 amino acids) and only slightly larger N-termini (median=97 amino acids) and most (95%) do not contain any predictable domain in addition to the catalytic domain. Of the 13 ALPHs that have a further domain, three have domains with functions in cytochrome c complex assembly (IPR021150, IPR018793) indicating mitochondrial functions and five ALPHs have a THIF-type NAD/FAD binding fold, usually found in the ubiquitin activating E1 family, indicating a possible function in protein degradation. ALPH of *Lentinula edodes* has a Peroxin-3 domain, indicating a peroxisomal function. Two ALPHs of *Rachicladosporium* have Pectate lyase domains, indicating a possible function degradation of cell wall material. ALPH of *Rhizopogon vesiculosus* has a second ALPH domain.

**Figure 3:**
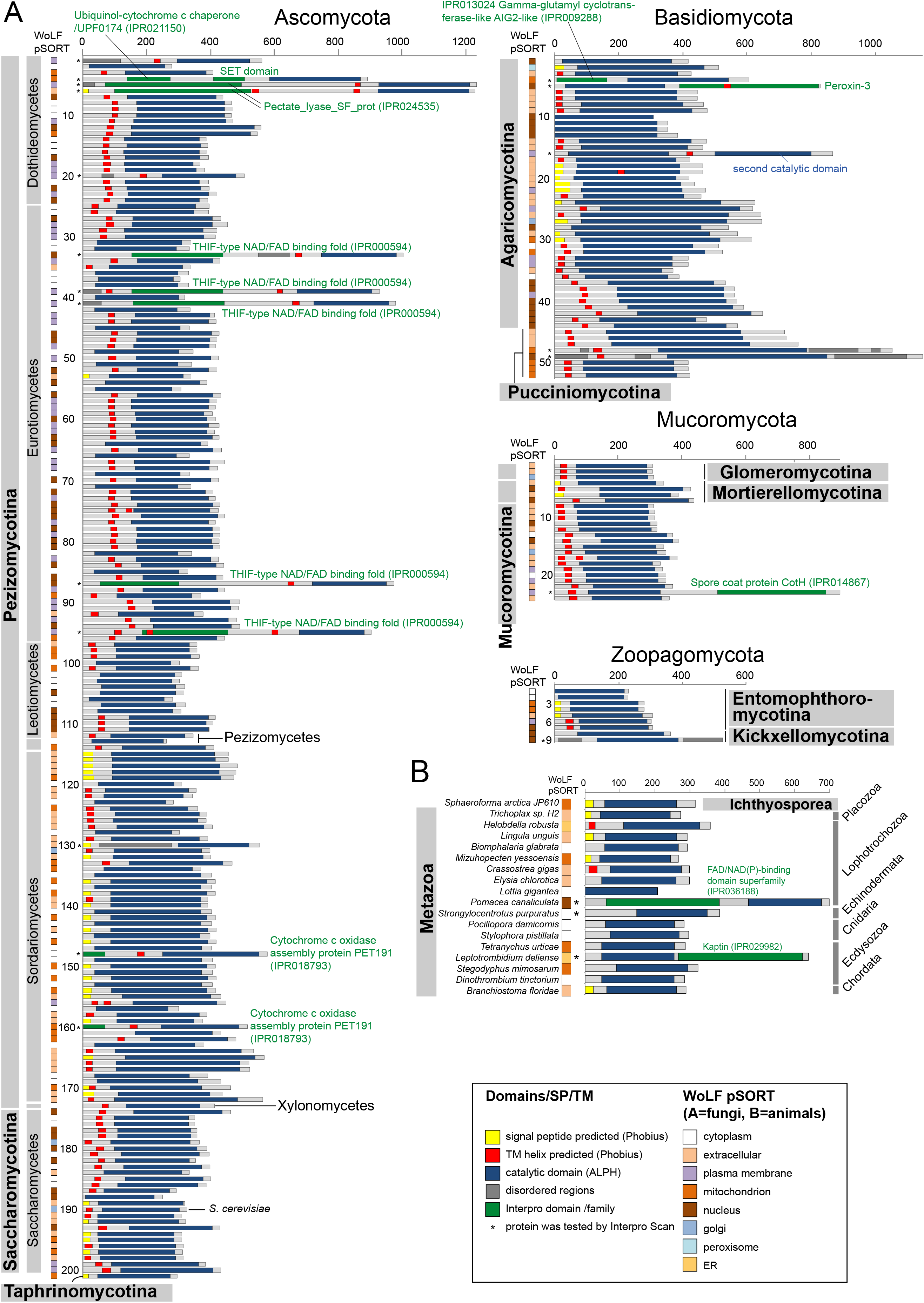
ALPHs of Opisthokonts. ALPHs of Dikarya **(A)** and ALPHs of Holozoa **(B)** are presented with their catalytic domain (dark blue) as well as further predicted sequence features, domains and localisation predictions. The organism names for the Dikarya ALPHs can be found in Supplementary Table S1c (number). More details on both Dikarya and Holozoa ALPHs are listed in Supplementary Table S1c and S1d, respectively.

**Figure 4:**
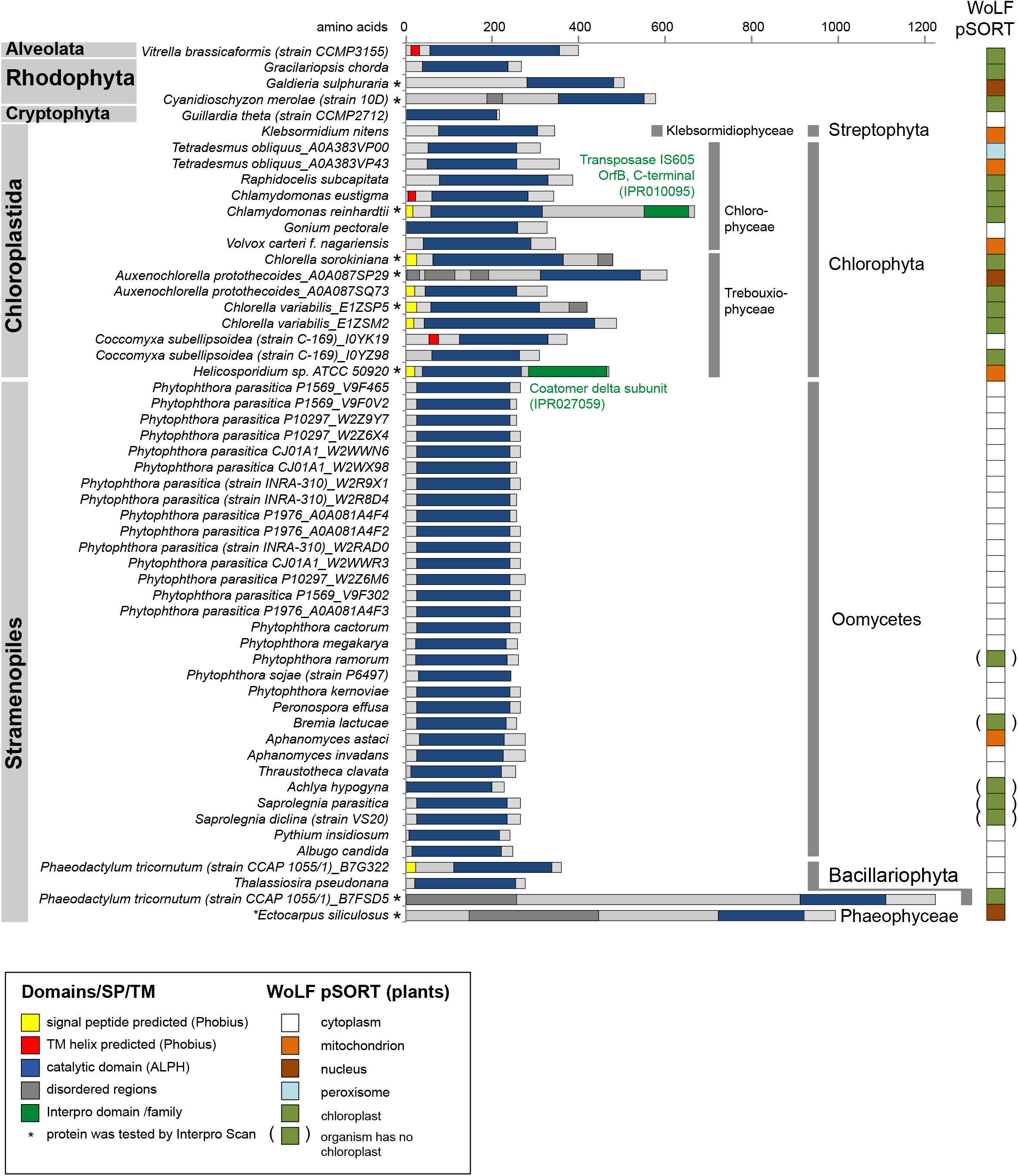
ALPHs of Diaphoretickes. ALPHs of Diaphoretickes are presented with their catalytic domain (dark blue), further predicted sequence features and domains and localisation predictions. More details are listed in Supplementary Table S1e.

The most interesting finding was the presence of a predicted trans-membrane region and/or signal peptide within the C-terminal region in 79% of all Dikaryan ALPH proteins. 184 ALPH proteins have predicted membrane helices (mostly one), a further 44 ALPH proteins have predicted signal peptides and only 61 ALPH proteins have neither predicted (Fig. 3A, Table S1c). In agreement with these data, only 63 ALPH proteins have predicted cytoplasmic localisation (mostly the ones without predicted membrane helices and signal peptides). The remaining proteins are predicted to be extracellular (31.8%), in the nucleus (27.8%), at the plasma membrane (21.1%), in the mitochondrion (16.1), in the golgi (2.7%) or in peroxisomes (0.4%). Prediction data need to be considered with care in the absence of experimental evidence, but taken together, the data provide strong evidence for most Dikaryan APLH proteins being non-cytoplasmic. Experimental data confirm non-cytoplasmic localisation for the ALPH protein of *S. cerevisiae* (YNL217w, Ppn2): two high-throughput studies indicate vacuolar localisation [35,36] and recent data show that Ppn2 is delivered to the vacuolar lumen via the multivesicular body pathway, where it functions as an endopolyphosphatase [7].

ALPHs are underrepresented in Holozoans (Fig. 1). In particular, all 130 vertebrate proteomes lack ALPHs and of the three available non-vertebrate proteomes of Chordata, only the Lancelet *Branchiostoma floridae* is ALPH-positive. Only four of 140 available Ecdysozoan proteomes contain ALPH; all are Arachnida. ALPHs are present in subgroups of Cnidarians (2/3), Echinodermatans (1/2), Lophotrochozoens (8/11), Placozoans (1/2) and Ichthyosporeans (1/2). 7 of these 18 Holozoan ALPH proteins have a predicted signal peptide or transmembrane helix at their C-termini and a non-cytoplasmic localisation prediction (mitochondrion, extracellular or ER); an additional 5 proteins have a predicted non-cytoplasmic localisation only (Fig. 3B, supplementary table S1d). Two proteins have an additional domain: ALPH of *Pomacea canaliculata* has a domain of the FAD/NAD(P)-binding superfamily N-terminal of the catalytic domain and ALPH of *Leptotrombidium deliense* may be interacting with actin, as it has a Kaptin domain C-terminal of its ALPH domain (Fig. 3B, and Supplementary Table S1d).

### ALPHs of Diaphoretickes

We found no ALPHs in land plants (100 proteomes), Apicomplexa (20 proteomes) or Ciliata (3 proteomes). ALPH is present in 3/4 species of red algae, 11/18 species of green algae (Chlorophyta), in the filamentous green algae *Klebsormidium nitens*, in the photosynthetic Alveolate *Vitrella brassicaformi* and in the cryptophyte algae *Guillardia theta.* 23/28 Stramenopiles have ALPHs: These are mostly (non-photosynthetic) Oomycetes, including all strains of *Phytophthora parasitica*; these plant pathogens have with 3 ALPH isoforms an unusually large number. Predicted signal peptides or transmembrane helices are present at the C-termini of many Chloroplastida ALPHs, ALPHs of *Vitrella brassicaformi* and in the ALPH of the diatom *Phaeodactylum tricornutum;* in many cases this correlated with a predicted localisation to the chloroplast. In contrast, ALPHs of Oomycetes have very short N- and C-termini and mostly a predicted cytoplasmic localisation. Of all 55 ALPHs of Diaphoretickes, only two have additional domains: *Chlamydomonas reinhardtii* ALPH has a predicted Transposase IS605 domain (a transposon of bacterial origin) and ALPH of *Helicosporidium* has a coatomer delta subunit domain.

### ALPHs of Euglenozoa

ALPHs are present in all Kinetoplastida and in their close relative *Euglena gracilis* (Supplementary Table Sf). The only possible exception is the free-living, non-parasitic *Bodo Saltans*, but this genome is not yet complete.

ALPHs of Kinetoplastida fall into two groups: Each organism has exactly one ALPH that is homologous to *T. brucei* ALPH1, the mRNA decapping enzyme [10]. These ALPHs all have a C-terminal extension of between 220 and 278 nucleotides and, with two exceptions (*Leptomonas pyrrhocoris* and *T. grayi*), they all have N-terminal extensions of a similar size. The *in vitro* mRNA decapping activity of *T. brucei* ALPH1 does not require the N-terminus [10], and the two ALPHs that lack the N-terminus are therefore likely active in mRNA decapping too. The fact that no Kinetoplastida strain has lost its ALPH1 homologue, and the absence of homologues to the canonical mRNA decapping enzymes (DCP1/DCP2), indicates that all Kinetoplastida rely on ALPH1 for mRNA decapping. In addition to the ALPH1 homologue, some *Leishmania* strains and all *Trypanosoma* and *Paratrypanosoma* have between one and three additional ALPH proteins that consist exclusively of the catalytic domain; four Leishmania strains have extensions within this domain. The absence of non-ALPH1 homologues in some *Leishmania* strains as well as in *Leptomonas*, *Crithidia* and *Blechomonas* indicates a less general and perhaps less essential function of these ALPH proteins.

When a phylogenetic tree is constructed from the sequences of the catalytic domains, the Kinetoplastida ALPH1 homologues form a separate group, distant to the group of the non-ALPH1 homologues, indicating that the ALPH1 decapping enzyme has evolved only ones in the last common ancestor of the Kinetoplastida (Figure 5A).

**Figure 5:**
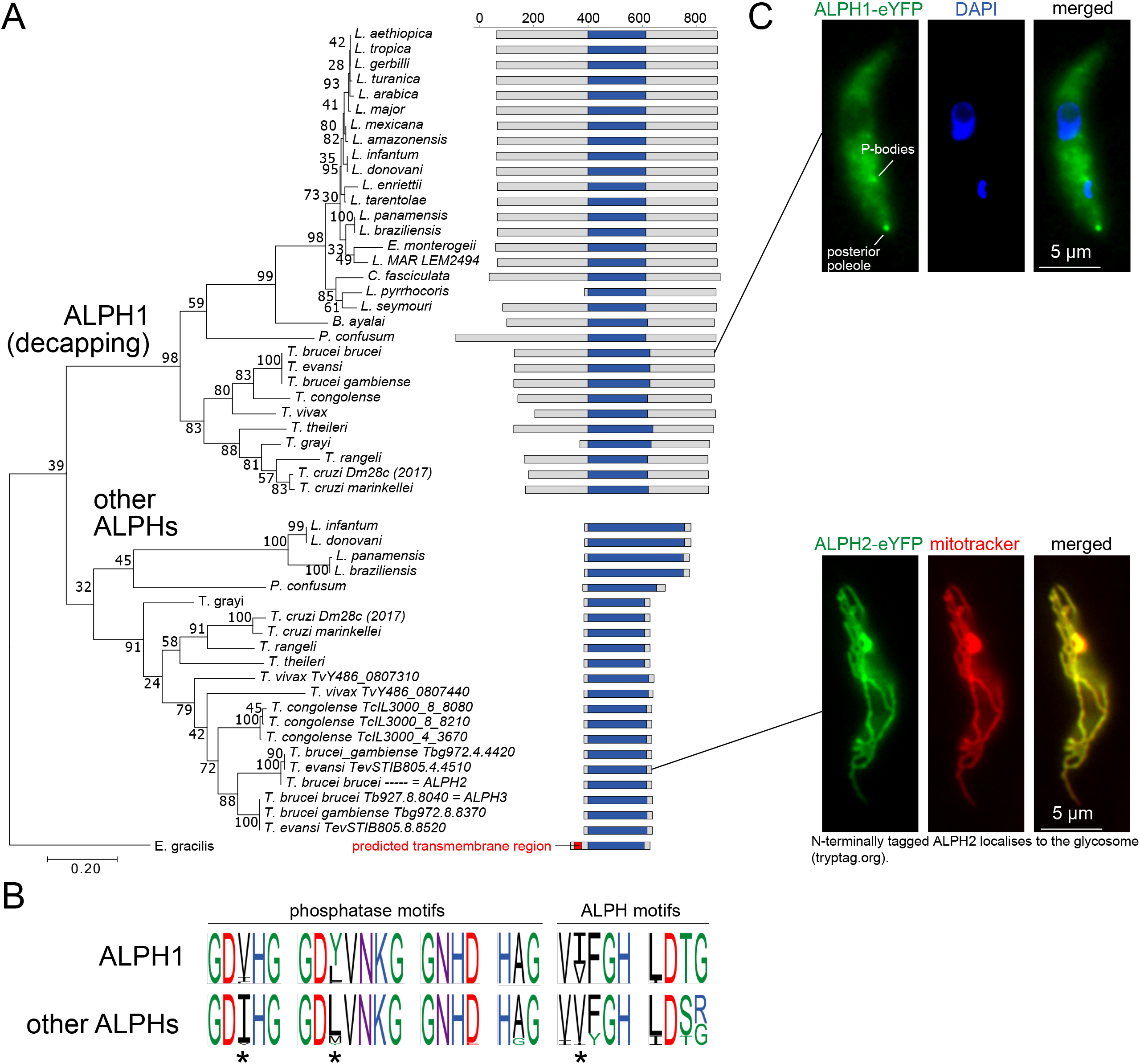
ALPHs of *Euglenozoa*. **(A)** ALPHs of *Euglenozoa* are presented with their catalytic domain (dark blue) along a maximum likelihood phylogenetic tree based on an alignment of gap corrected sequences of ALPH catalytic domains. Different methods (minimal evolution, neighbour joining) gave slightly different trees, but with all methods, the ALPH1 isoforms group together in one clade that never contains non-ALPH1 isoforms. All details on *Euglenozoa* ALPHs are listed in Table S1f. The localisation of *T. brucei* ALPH1 and ALPH2 was determined by expressing eYFP fusion proteins. One representative fluorescent image of each cell line is shown. At least three different clonal cell lines showed identical localisation. **(B)** Sequence motifs of the ALPH1 orthologues and the Kinetoplastida orthologues to other ALPHs. Major differences are marked by asterisks.

None of the ALPH sequences from Kinetoplastida contains any recognisable domain or localisation signal. The decapping enzyme *T. brucei* ALPH1, fused to eYFP either C- or N terminally, localises to the cytoplasm, P-bodies and to the posterior pole of the cell and this cytoplasmic localisation is consistent with the essential function of ALPH1 in mRNA decapping ([10] Figure 5A). To investigate the localisation of an non-ALPH1 homologue, *T. brucei* ALPH2 was expressed as C-terminal eYFP fusion in procyclic trypanosomes using an inducible expression system [37]. ALPH2 showed a localisation pattern characteristic of mitochondrial proteins and co-localised with a mitochondrial stain (MitoTracker^TM^ Orange) (Figure 5A).

Differences between the two different groups of Kinetoplastida ALPH proteins are also obvious within the phosphatase and ALPH motifs: three positions show major differences; most pronounced is the preference of GDVHG in decapping enzymes and GDIHG in the non-decapping ALPH proteins.

Euglena is too distantly related to the Kinetoplastida to unequivocally design its ALPH to the ALPH1 or non ALPH1 group, but its first and second sequence motif (GDIHG and GDLVGKG) indicates a non-ALPH1, and this is also suggested by the absence of N- and C-terminal sequences (Figure 5B). Interestingly, Euglena ALPH is the only Kinetoplastida ALPH with a predicted trans-membrane domain. Euglena ALPH was not enriched within purified mitochondria fractions [38] and also not within chloroplasts (Martin Zoltner, Charles University in Prague, personal communication). Like Kinetoplastida, Euglena has no recognisable homologue to the canonical mRNA decapping enzyme DCP1/DCP2 in its genome the absence of an ALPH1 homologue raises the question of how mRNA decapping is achieved in this organism.

### ALPHs have *in vitro* mRNAs decapping activity

The phylogeny data from us and others [6] show that ApaH like phosphatases were present in the last common ancestor of eukaryotes. Since then, large percentages of eukaryotes have lost the enzyme and of those ALPHs that are still present, large percentages have predicted non-cytoplasmic localisations. The finding that the trypanosome ALPH1 evolved to be the parasites mRNA decapping enzyme [10] prompted us to test, whether ALPHs from other organisms have mRNA decapping activity too. If true, this could provide an explanation for the evolutionary thrive against the presence of cytoplasmic ALPHs in eukaryotes, concurrent with the evolvement of the eukaryotic mRNA cap.

We produced recombinant ALPH proteins from randomly selected organisms of the all three eukaryotic kingdoms that have ALPHs: we used ALPH of the Ichthyosporea *Sphaeroforma arctica,* ALPH of the green algae *Auxenochlorella protothecoides* and ALPH2 from *Trypanosoma brucei* (ALPH2 has mitochondrial localisation and is not the mRNA decapping enzyme). As a control, ALPH of *Sphaeroforma arctica* was also produced as an inactive mutant by mutating a conserved amino acid in the metal ion binding motif (GDVIG:GNVIG [39]). All four proteins were purified from Arctic express cells (Supplementary Figure S2A) and subsequently tested in *in vitro* decapping assays using a 39 nucleotide long RNA oligo with a m^7^G cap structure as a substrate. Capped and uncapped oligos can be distinguished by differences in gel mobility on urea acrylamide gels complemented with acryloylaminophenyl boronic acid [40,41]. To identify the optimal reaction conditions for each enzyme, the decapping activity of the three active enzymes was first tested in the presence of different bivalent ions (Mg^2+^, Mn^2+^, Co^2+^ and Zn^2+^) (Supplementary Figure S2B). All enzymes had mRNA decapping activity, but the ion requirements for optimal activity differed: ALPH of *Sphaeroforma arctica* showed highest decapping activity with Mn^2+^ and some activity with Co^2+^, but none with Mg^2+^ or Zn^2+^. *T. brucei* ALPH2 had best decapping activity with Co^2+^ and some with Mg^2+^ and possibly Mn^2+^, but none with Zn^2+^. *Auxenochlorella protothecoides* ALPH had best activity with Mg^2+^ and some with Co^2+^, but none with Zn^2+^ and Mn^2+^. Figure 6 shows the results of one representative decapping assay for each of the enzymes, in the presence of the optimal ion, and, as a control with one ion that promotes enzyme activity not or little. Importantly, the catalytically inactive mutant ALPH of *Sphaeroforma arctica* had no decapping activity. Together with the differences in ion requirements, this is strong evidence that the activity of the wild type proteins is not caused by a contaminating bacterial enzyme. The *T. brucei* decapping enzyme ALPH1 served as a positive control: it has highest activity with Mg^2+^ and reduced activity with Mn^2+^. The data show that all ALPH enzymes tested accept capped mRNA as a substrate *in vitro*.

**Figure 6:**
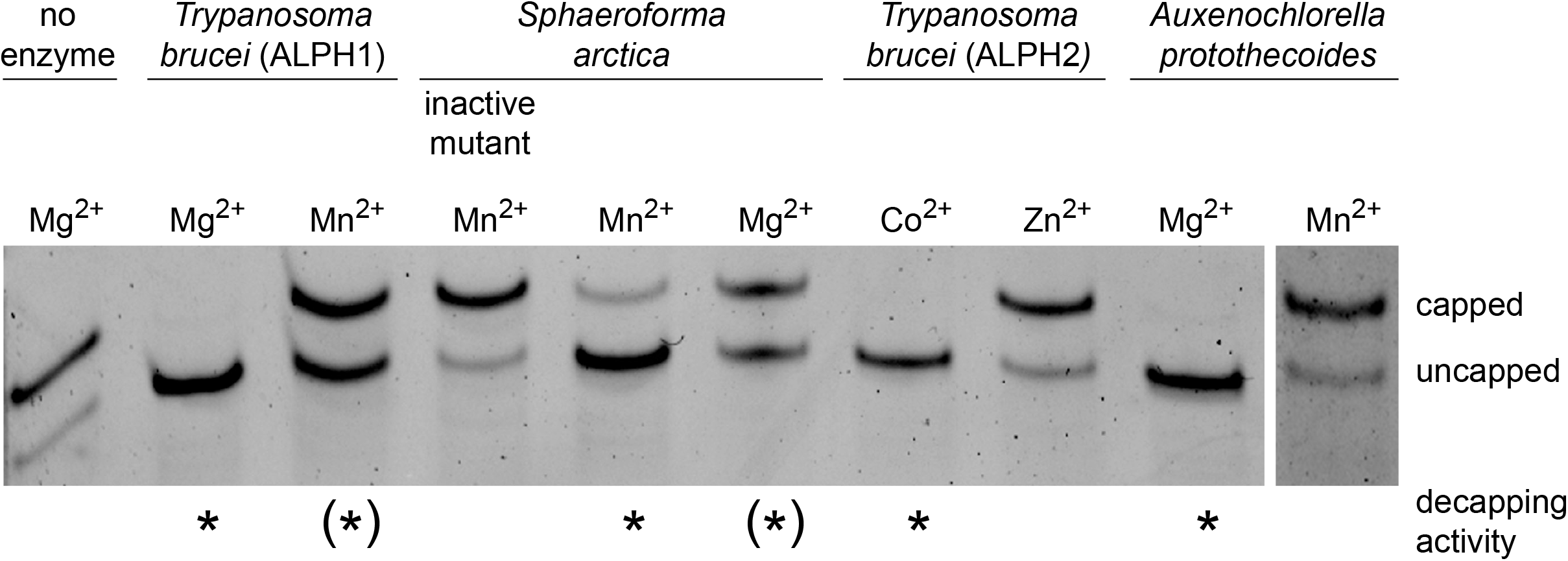
ALPH proteins have mRNA decapping activity. *In vitro* mRNA decapping assays were done with recombinant ALPH enzymes from *T. brucei* (the decapping enzyme ALPH1 and the mitochondrial localised enzyme ALPH2), from *Sphaeroforma arctica* (both the wild type and a catalytically inactive mutant) and from *Auxenochlorella protothecoides,* using an m^7^G-capped RNA oligo as a substrate. Note that a small portion of this oligo is uncapped even in the absence of enzyme due to production issues (there is no auto-decapping activity during the assay, data not shown). mRNA decapping activity was observed, as expected, with *T. brucei* ALPH1 [10] and was fully absent in the catalytically inactive mutant of ALPH from *Sphaeroforma arctica*. All three previously untested ALPH proteins had *in vitro* decapping activity, albeit the ions requirements differed between the enzymes.

## DISCUSSION

mRNAs are essential molecules in both eukaryotes and prokaryotes and need to be protected from unregulated exonucleolytic degradation. 5’-3’exoribonucleases require monophosphorylated RNA 5’ends and the major protection strategy is therefore to avoid monophosphates at the mRNAs 5’end. Most eukaryotic mRNAs are modified at their 5’end by an m^7^ methylguanosine (m^7^G) cap, linked via an inverted 5’-5’bridge [42]. A small fraction of eukaryotic RNAs carries non-canonical nucleotide metabolite caps, for example an NAD+ cap [43,44]. Bacteria were long believed to simply keep the triphosphate that is naturally present at their 5’end after transcription, but more recently were found to have diphosphate 5’ends [45], and also non-canonical metabolite caps, such as the NAD+ cap [46], the 3-dephospho-coenzyme A cap (dpCoA) [47] or dinucleoside polyphosphate caps [24–26]. Archaea mRNAs are not well studied by do carry triphosphates caps [48].

In common to the increasing number of different mRNA caps and mRNA 5’ end modifications discovered throughout all kingdoms of life is the pyrophosphate bond, and therefore the requirement of a pyrophosphatase to remove the cap or the polyphosphate. Enzymes of four different families can cleave pyrophosphate bonds at the 5’end of mRNAs [49]. DXO and histidine triad (HIT) proteins, act mainly on faulty RNAs or on cap remnants or remove non-canonical NAD+ caps without cleaving the pyrophosphate bond [49]. This leaves two pyrophosphatase families that act on intact mRNAs. Nudix domain proteins include the prototypic eukaryotic mRNA decapping enzyme Dcp2 [50] and several further eukaryotic nudix hydrolases with activity to the m^7^G cap and non-canonical metabolite caps [51–53] as well as the bacterial RppH [54] and NudC proteins [55] that act on di- or triphosphorylated mRNAs and on NAD+ caps [49]. Enzymes of the fourth group, the ApaH family only entered the field of mRNA decapping most recently. ApaH like phosphatase ALPH1 from *Trypanosoma brucei* was shown to be the only or major mRNA decapping enzyme in these parasites [10]. Bacterial ApaH cleaves of non-canonical nucleoside-tetraphosphate caps [24–26] and likely regulates gene expression during stress.

The major aim of this work was to find out, how widespread mRNA decapping by ApaH like phosphatases is. To identify ALPH proteins in all available eukaryotic proteomes, we have developed a Python based algorithm based on recognising the six conserved motifs of ALPH. We have started with a well-defined training set [6] and stepwise optimised the matrix of the motifs to recognise all ALPHs, but no related phosphatases. Our comprehensive datasets on the 412 ALPH proteins are summarized in detail in Table S1 and contribute to a more comprehensive knowledge of this enzyme family.

Our data provide no evidence for the presence of additional mRNA decapping enzymes among eukaryotic ApaH-like phosphatases. We expected putative mRNA decapping enzymes to fulfil two criteria: they must be cytoplasmic and they must possess additional sequences next to the catalytic domain that can regulate enzyme activity. The *Trypanosoma brucei* decapping enzyme ALPH1 is cytoplasmic and has C- and N-terminal extensions. Outside the Kinetoplastida, most ALPHs consist of the catalytic domain only and have predicted non-cytoplasmic localisations, often further supported by predicted transmembrane helices and signal peptides. In the few cases when additional domains were present, these did not indicate any connection to mRNA metabolism or showed any similarities to the Kinetoplastida C- and N-terminal extensions.

Surprisingly, all three ALPH proteins that we tested, had mRNA decapping activity *in vitro*, even though none of these enzymes is likely to use mRNA as their physiological substrate: *T. brucei* ALPH2 localises to the mitochondrion (Figure 5A) and ALPHs of *Sphaeroforma arctica* and *Auxenochlorella protothecoides* both have predicted signal peptides and predicted localisation to the mitochondrion and chloroplast, respectively. This rather broad substrate range of ALPHs is not unexpected: the related bacterial ApaH protein cleaves NpnN nucleotides [13–15] as well was mRNA capped with NpnN [24,26] and even has phosphatase and ATPase activity in addition to its pyrophosphatase activity [56]. The only other characterised ALPH protein, the *S. cerevisiae* ALPH protein Ppn2 has, to our knowledge, not been tested for mRNA decapping activity. Substrate unspecificity is not restricted to ApaH and ApaH like phosphatases, but is also found in enzymes of the nudix domain family [57]: the major bacteria decapping enzyme RppH was initially identified as diadenosine polyphosphate hydrolase [58] and can also cleave NAD+ caps [53] and recent in vitro studies of mammalian nudix proteins shows that many have mRNA decapping activity towards a range of caps and some also towards free metabolites [51,53,59,60]; how many of these nudix proteins have mRNAs as their natural substrates is still debated.

The broad substrate range of ALPHs that includes mRNA caps provides an explanation for its loss or non-cytoplasmic localisation in most eukaryotes. The last common ancestor of all eukaryotes had ALPH. This assumption is supported by the presence of ALPH proteins in all eukaryotic kingdoms found in this and earlier [6] studies. With the evolvement of the eukaryotic mRNA cap, eukaryotes faced two problems: first, the cap needed to be protected from unregulated degradation by pyrophosphatases and second, an mRNA decapping enzyme was needed for regulated degradation. The first challenge was likely addressed in multiple ways, including for example protection by cap binding proteins like the eIF4G complex. However, one important evolutionary strategy may have been to inactivate the bacterial-inherited ALPH proteins. Many eukaryotes lost ALPH proteins, others have transferred them into organelles away from mRNA substrates, where they fulfil essential functions, for example in poly-phosphate metabolism [7]. Evolution has solved the second challenge by transforming available pyrophosphatases with wide substrate ranges into regulatable mRNA decapping enzymes. Mostly nudix hydrolases were exploited for this task, but Kinetoplastida uniquely use one of their ALPH proteins.

One open question is, whether all organisms outside the Kinetoplastida use DCP2 orthologues as their major mRNA decapping enzyme. While DCP2 has been functionally characterised in yeast [50], human [61] and plants [62], there are no experimental data on Metamonada and Discoba, and DCP2 orthologues are difficult to identify by blast searches alone. Metamonada possibly have a DCP2 homologue: in *Trichomonas* this can be identified by Blast and in *Giardia* by more sophisticated *in silico* methods [63]. Whether Discoba have DCP2 is unclear. A possible DCP2 homologue is identifiable by blast in the Discoba *Naegleria*, but not in Euglena. The question whether DCP2 was present in the last common ancestor and selectively lost in Kinetoplastida, or whether it evolved subsequently to the separation of the Discoba cannot be answered without experimental data.

In the last few years, more and more mRNA cap variants have been discovered in both prokaryotes and eukaryotes, and more and more enzymes that act in mRNA decapping. Our findings that ApaH like phosphatases have mRNA decapping activity, but, with the exception of the Kinetoplastida enzymes, are not involved in mRNA decapping in eukaryotes is an important contribution towards a more comprehensive picture of mRNA decapping.

## CONCLUSIONS

Despite of being present in all eukaryotic kingdoms, ApaH like phosphatases are poorly studied. Here, we present a new algorithm for unequivocal classification of a protein as an ALPH and we provide a comprehensive dataset of ApaH like phosphatases of all available reference proteomes (Table S1). 500/824 proteomes had no ALPH and the remaining 324 proteomes contained, in total, 412 ALPH proteins.

Almost all ALPHs consisted of the catalytic domain only and had non-cytoplasmic localisation predictions supported by the presence of transmembrane helices and signal peptides. The marked exceptions are ALPHs of the Kinetoplastida that are orthologues to the *Trypanosoma brucei* mRNA decapping enzyme ALPH1: all have C- and most have N-terminal extensions. Despite of their non-cytoplasmic localisations that excluded mRNAs as their natural substrates, we found that ALPH proteins from all eukaryotic kingdoms had mRNA decapping activity *in vitro*, suggesting that (i) the loss or non-cytoplasmic localisation of ALPHs in most eukaryotes was caused by evolutionary pressure to protect mRNAs from uncontrolled degradation and (ii) the N- and C-terminal extensions of the Kinetoplastida ALPH proteins serve to regulate the mRNA decapping activity of ALPH. mRNA decapping is one of only two known ALPH functions, but our data strongly suggest that this function is restricted to ALPHs of Kinetoplastida. The physiological functions of most other ALPHs remain to be discovered. *In vivo* experiments are essential, as the wide substrate range of ALPH will impede the identification of physiological substrates *in vitro*.

## METHODS

### Identification of ApaH like phosphatases

A novel algorithm was developed using Python 3 to identify ALPHs in a set of proteome fasta files. The algorithm initially used narrow matrices to identify the 6 conserved motifs (4 common to all PPP and 5-6 unique to ALPHs) based on the data of [3,6], these matrices were successively extended after training on a set of 46 yeast proteomes, controlled by BLAST, and later used to screen 824 eukaryotic proteomes. This screen gave information about conserved and non-conserved distances between the 6 motifs. All proteins that had only one motif missing were manually inspected further. If they possessed the missing motif in an only slightly, conservatively-altered version and, at the same time, had the correct distances between the motifs (where these distances are conserved), the motif matrix was extended to include this motif. The final version of the algorithm uses one narrow matrix for the initial screen: motive 1 [“G”, “D”, “VILT”, “HQ”, “G”]; motive 2 [“G”, “DN”, “LMIFVT”, “IVTLEAC”, “NSATFGVEQIDHM”, “KQN”, “GASHT”]; motive 3 [“G”, “N”, “HNWQD”, “ED”]; motive 4= [“H”, “AG”, “G”]; motive 5 [“VIT”, “IVYFLAMT”, “FY”, “G”, “H”]; motive 6 [“LVIMT”, “DE”, “STGL”, “GASRN”]. If not all motifs were found, the algorithm tolerates up to two motives from an extended matrix: motive 1 [“GxX”, “DPEKRxX”, “QWERTIPASDFGHKLYCVNMxX”, “HAQxX”, “GFSAxX”]; motive 2 [“GxX”, “DNxX”, “QWERTIPASDFGHKLYCVNMxX”, “ILCVTMEAxX”, “NDSTAGCHMFEVQIxX”, “KQNxX”, “GASHTxX”]; motive 3 [“GxX”, “NxX”, “HNTWQDxX”, “EDxX”]; motive 4 [“HxX”, “AGLVxX”, “GxX”]; motive 5 [“VILCMTxX”, “QWERTIPASDFGHKLYCVNMxX”, “FYCxX”, “GxX”, “HxX”]; motive 6 [“LVIMTGxX”, “DExX”, “ATGSVLxX”, “GRASNExX”]. Furthermore, the programme restricts the distances between the motifs to 14-90 (motive 1 to 2), 25-80 (motive 2 to 3) and 17-30 (motive 5 to 6). ALPHs with missing motives due to wrong-annotated start codons or sequencing errors will not be detected, with the exception of an unidentified amino acid included in the extended matrix (x, X). Only for the Kinetoplastida, such ALPHs were manually included. Initially, the software tolerated R at position 6 in motive 2. However, distinction between ALPHs and ApaH / RLPH was not possible in all cases and therefore R at this position was not tolerated by either matrix any longer. This affected 19 proteins that had been previously recognised as an ALPH (14 discoba, 3 metazoa and 2 embryophyta). The Python programme is available upon request.

### Cloning and expression of recombinant proteins

The mRNA decapping enzyme *T. brucei* ALPH1 was expressed as an N-terminal truncation with a C-terminal 6xHis-Tag in RosettaBlue competent bacteria cells (Novagen) as previously described [10].

All other proteins were expressed either full length (*T. brucei* ALPH2 (Tb927.4.4330)) or with minor N-terminal truncations to exclude the transmembrane regions *(Sphaeroforma arctica:* amino acids 21-316; *Auxenochlorella protothecoides:* amino acids 21-327) in Arctic Express DE competent cells (Agilent). At the N-terminus, the proteins were fused to a tag consisting of a 6xHis-Tag, a Thrombin cleavage site, a SUMO tag and a TEV cleavage site, resulting in the following additional protein sequence:

MGSSHHHHHHSSGLVPRGSASMSDSEVNQEAKPEVKPEVKPETHINLKVSDGSSEIFFKIKKTTPLRRLMEAFAKRQGKEMDSLRFLYDGIRIQADQTPEDLDMEDNDIIEAHREQIGGHMENLYFQGEASAT. At the C-terminus, all proteins had additional GSGSGC, due to cloning reasons. For the inactive mutant of the *Sphaeroforma arctica* ALPH, the highly conserved GDVIG motif involved in metal ion binding was mutated to GNVIG [39]. The purification of the recombinant proteins was done via Nickel-agarose beads using standard procedures as previously described [10], except that expression in Arctic Express DE cells was done at 10°C, following the instructions of the company. The concentration of all purified proteins was estimated from Coomassie gels using a titration of a protein with known concentration for calibration.

### Phylogeny

The Maximum likelihood tree of the Euglenozoa ALPHs (Figure 5) was done with all ALPH catalytic domains (defined as starting 6 amino acids upstream of motif 1 and ending 12 amino acids downstream of motif 6) using MEGA 7 [64] with 500 bootstrap cycles. The sequences of *Crithidia fasciculata* and *Trypanosoma vivax* were gap-corrected: one gap was removed).

### Proteomes

Proteomes were downloaded from Uniprot, using all available reference proteomes of UniProt_release 2019_02 (For Archaea: 2019_11). [27]. Table S1a lists all proteomes used in this work. For Kinetoplastida, we downloaded all available proteomes from TriTrypDB (February 2019) [28,29].

### Work with trypanosomes

*Trypanosoma brucei* Lister 427 procyclic cells expressing a TET repressor (pSRP2.1) were used for all experiments [37]. Cells were cultured in SDM-79 [65] at 27 °C and 5% CO_2_. Transgenic trypanosomes were generated using standard procedures [66]. All experiments used logarithmically growing trypanosomes. The mitochondria were stained by adding 1 μM MitoTracker™ Orange CMTMRos (Invitrogen) to living cells; imaging was done within 30 minutes.

### Microscopy

Z-stack images (100 stacks at 100 nm distance) were taken with a custom build TILL Photonics iMic microscope equipped with a sensicam camera (PCO), deconvolved using Huygens Essential software (Scientific Volume Imaging B. V., Hilversum, The Netherlands). All images are presented as Z-projections (method sum sliced) produced by ImageJ software.

### *In vitro* decapping assays

Each decapping reaction was done in 10 μl volume (with RNAse-free water) containing 50 mM Tris-HCl (pH 7.9), 100 mM NaCl, 1 mM metal ions (variable), 40 units Ribolock RNAse inhibitor (ThermoFisher Scientific), 0.25 μM recombinantly-produced ALPH enzyme and 3.8 μM capped RNA oligo m^7^G-AACUAACGCUAUUAUUAGAACAGUUUCUGUACUAUAUUG (Bio-Synthesis, Texas, USA). The sample was incubated for 1 hour at 37°C. Control reactions were done without enzyme. The reaction was stopped by ethanol precipitation: 1 μl 3 M RNAse-free sodium acetate (pH 5.5) (Ambion), 1 μl RNAse-free glycogen (ThermoFisher Scientific) and 30 μl cold 100% ethanol was added, the sample incubated at −20°C for 30 minutes, followed by centrifugation (15 minutes, 20,000 g). The pellet (containing the RNA oligo) was washed once in 80% ethanol, air-dried, resolved in 5 μl RNAse-free water (ThermoFisher Scientific) and either stored at −80°C or directly prepared for loading to the gel.

RNA Samples were separated on urea acrylamide gels that contained acryloylaminophenyl boronic acid [40,41]: boronyl groups form stable complexes with the m^7^G cap structure this way increasing the difference in gel mobility between capped and uncapped oligos [40,41]. For 12% gels, a 10 ml gel solution was freshly prepared with distilled water, containing 12% Acrylamide/Bis-acrylamide (19:1) (Sigma-Aldrich), 7M urea (Sigma-Aldrich), 0.1% ammonium persulfate, 4 μl TEMED, 0.25% acryloylaminophenyl boronic acid (Sigma-Aldrich) and 100 mM Tris-acetate (pH 9.0). Gels were run in 0.5x TBE buffer (45 mM Tris-HCl pH 8.3, 45 mM boric acid, 1 mM EDTA) using the Mini-PROTEAN Tetra Vertical Electrophoresis cell (gel size 10×8 cm, BioRad). Each run was preceded by a 30 min pre-run at 300 V to warm the gel. The purified RNA samples were prepared for gel electrophoresis by adding 1 μl loading dye (gel loading buffer II, Invitrogen) followed by incubation at 95°C for 5 minutes. Samples were loaded immediately to the pre-rinsed wells of the pre-run gel and the gel was run at 200 V (after a 10 min run at 170 V) for 30-40 minutes, stained for 20 min in SYBR™ Green II RNA Gel Stain (1:10,000 in 0.5x TBE, ThermoFisher Scientific) and imaged on the iBright™ (Invitrogen) with nucleic acid settings.

## DECLARATIONS

### Ethics approval and consent to participate

Not applicable

### Consent for publication

Not applicable

### Availability of data

All data generated or analysed during this study are included in this published article [and its supplementary information files].

### Authors’ contributions

SK wrote the manuscript and did the *in silico* work of this study, with the exception of the phylogenetic tree of Figure S1 that was done by BB. PACL and NB did all wet work (protein expression, decapping assays).

## Acknowledgement

We like to thank Martin Zoltner (Charles University in Prague, Czech Republic), Mark Carrington (University of Cambridge, UK) and Mark Field (University of Dundee, UK) for sharing unpublished proteomes and data on Euglena and *Bodo Saltans*.

## REFERENCES

1. Kerk D, Templeton G, Moorhead GBG. Evolutionary radiation pattern of novel protein phosphatases revealed by analysis of protein data from the completely sequenced genomes of humans, green algae, and higher plants. Plant Physiol. American Society of Plant Biologists; 2008;146:351–67.

2. Shi Y. Serine/threonine phosphatases: mechanism through structure. Cell. 2009;139:468–84.

3. Andreeva AV, Kutuzov MA. Widespread presence of “bacterial-like” PPP phosphatases in eukaryotes. BMC Evol Biol. 2004;4:47.

4. Uhrig RG, Moorhead GB. Two ancient bacterial-like PPP family phosphatases from Arabidopsis are highly conserved plant proteins that possess unique properties. Plant Physiol. American Society of Plant Biologists; 2011;157:1778–92.

5. Uhrig RG, Labandera A-M, Moorhead GB. Arabidopsis PPP family of serine/threonine protein phosphatases: many targets but few engines. Trends Plant Sci. 2013;18:505–13.

6. Uhrig RG, Kerk D, Moorhead GB. Evolution of bacterial-like phosphoprotein phosphatases in photosynthetic eukaryotes features ancestral mitochondrial or archaeal origin and possible lateral gene transfer. Plant Physiol. 2013;163:1829–43.

7. Gerasimaitė R, Mayer A. Ppn2, a novel Zn2+-dependent polyphosphatase in the acidocalcisome-like yeast vacuole. J Cell Sci. 2017;130:1625–36.

8. Andreeva N, Ledova L, Ryazanova L, Tomashevsky A, Kulakovskaya T, Eldarov M. Ppn2 endopolyphosphatase overexpressed in Saccharomyces cerevisiae: Comparison with Ppn1, Ppx1, and Ddp1 polyphosphatases. Biochimie. Elsevier B.V; 2019;163:101–7.

9. Ryazanova LP, Ledova LA, Andreeva NA, Zvonarev AN, Eldarov MA, Kulakovskaya TV. Inorganic Polyphosphate and Physiological Properties of Saccharomyces cerevisiae Yeast Overexpressing Ppn2. Biochemistry Mosc. 2020;85:516–22.

10. Kramer S. The ApaH-like phosphatase TbALPH1 is the major mRNA decapping enzyme of trypanosomes. PLoS Pathog. 2017;13:e1006456.

11. Barton GJ, Cohen PT, Barford D. Conservation analysis and structure prediction of the protein serine/threonine phosphatases. Sequence similarity with diadenosine tetraphosphatase from Escherichia coli suggests homology to the protein phosphatases. Eur J Biochem. 1994;220:225–37.

12. Koonin EV. Bacterial and bacteriophage protein phosphatases. Mol Microbiol. 1993;8:785–6.

13. Plateau P, Fromant M, Brevet A, Gesquière A, Blanquet S. Catabolism of bis(5’-nucleosidyl) oligophosphates in Escherichia coli: metal requirements and substrate specificity of homogeneous diadenosine-5’,5’’’-P1,P4-tetraphosphate pyrophosphohydrolase. Biochemistry. 1985;24:914–22.

14. Guranowski A, Jakubowski H, Holler E. Catabolism of diadenosine 5’,5”‘-P1,P4-tetraphosphate in procaryotes. Purification and properties of diadenosine 5’,5”‘-P1,P4-tetraphosphate (symmetrical) pyrophosphohydrolase from Escherichia coli K12. J Biol Chem. 1983;258:14784–9.

15. Guranowski A. Specific and nonspecific enzymes involved in the catabolism of mononucleoside and dinucleoside polyphosphates. Pharmacol Ther. 2000;87:117–39.

16. Kimura Y, Tanaka C, Sasaki K, Sasaki M. High concentrations of intracellular Ap4A and/or Ap5A in developing Myxococcus xanthus cells inhibit sporulation. Microbiology (Reading). 2017;163:86–93.

17. Ismail TM, Hart CA, McLennan AG. Regulation of dinucleoside polyphosphate pools by the YgdP and ApaH hydrolases is essential for the ability of Salmonella enterica serovar typhimurium to invade cultured mammalian cells. J Biol Chem. American Society for Biochemistry and Molecular Biology; 2003;278:32602–7.

18. Hansen S, Lewis K, Vulić M. Role of global regulators and nucleotide metabolism in antibiotic tolerance in Escherichia coli. Antimicrob Agents Chemother. 2008;52:2718–26.

19. Nishimura A, Moriya S, Ukai H, Nagai K, Wachi M, Yamada Y. Diadenosine 5’,5’“-”P1,P4-tetraphosphate (Ap4A) controls the timing of cell division in Escherichia coli. Genes Cells. 1997;2:401–13.

20. Johnstone DB, Farr SB. AppppA binds to several proteins in Escherichia coli, including the heat shock and oxidative stress proteins DnaK, GroEL, E89, C45 and C40. EMBO J. 1991;10:3897–904.

21. Monds RD, Newell PD, Wagner JC, Schwartzman JA, Lu W, Rabinowitz JD, et al. Di-adenosine tetraphosphate (Ap4A) metabolism impacts biofilm formation by Pseudomonas fluorescens via modulation of c-di-GMP-dependent pathways. J. Bacteriol. 2010;192:3011–23.

22. Ji X, Zou J, Peng H, Stolle A-S, Xie R, Zhang H, et al. Alarmone Ap4A is elevated by aminoglycoside antibiotics and enhances their bactericidal activity. Proc Natl Acad Sci USA. 2019;116:9578–85.

23. Farr SB, Arnosti DN, Chamberlin MJ, Ames BN. An apaH mutation causes AppppA to accumulate and affects motility and catabolite repression in Escherichia coli. Proc Natl Acad Sci USA. 1989;86:5010–4.

24. Luciano DJ, Levenson-Palmer R, Belasco JG. Stresses that Raise Np4A Levels Induce Protective Nucleoside Tetraphosphate Capping of Bacterial RNA. Mol Cell. Elsevier Inc; 2019;:1–19.

25. Luciano DJ, Belasco JG. Np4A alarmones function in bacteria as precursors to RNA caps. Proc Natl Acad Sci USA. 2020;117:3560–7.

26. Hudeček O, Benoni R, Reyes-Gutierrez PE, Culka M, Šanderová H, Hubálek M, et al. Dinucleoside polyphosphates act as 5’-RNA caps in bacteria. Nat Commun. Nature Publishing Group; 2020; 11:1052–11.

27. UniProt Consortium. UniProt: a worldwide hub of protein knowledge. Nucleic Acids Res. 2019;47:D506–15.

28. Aslett M, Aurrecoechea C, Berriman M, Brestelli J, Brunk BP, Carrington M, et al. TriTrypDB: a functional genomic resource for the Trypanosomatidae. Nucleic Acids Res. 2010;38:D457–62.

29. Warrenfeltz S, Basenko EY, Crouch K, Harb OS, Kissinger JC, Roos DS, et al. EuPathDB: The Eukaryotic Pathogen Genomics Database Resource. Methods Mol Biol. 2018;1757:69–113.

30. Adl SM, Bass D, Lane CE, Lukes J, Schoch CL, Smirnov A, et al. Revisions to the Classification, Nomenclature, and Diversity of Eukaryotes. J Eukaryot Microbiol. John Wiley & Sons, Ltd (10.1111); 2018;66:jeu.12691-116.

31. Crooks GE, Hon G, Chandonia J-M, Brenner SE. WebLogo: a sequence logo generator. Genome Res. Cold Spring Harbor Lab; 2004;14:1188–90.

32. Mitchell AL, Attwood TK, Babbitt PC, Blum M, Bork P, Bridge A, et al. InterPro in 2019: improving coverage, classification and access to protein sequence annotations. Nucleic Acids Res. 2019;47:D351–60.

33. Käll L, Krogh A, Sonnhammer ELL. Advantages of combined transmembrane topology and signal peptide prediction--the Phobius web server. Nucleic Acids Res. 2007;35:W429–32.

34. Horton P, Park K-J, Obayashi T, Fujita N, Harada H, Adams-Collier CJ, et al. WoLF PSORT: protein localization predictor. Nucleic Acids Res. 2007;35:W585–7.

35. Huh W-K, Falvo JV, Gerke LC, Carroll AS, Howson RW, Weissman JS, et al. Global analysis of protein localization in budding yeast. Nature. 2003;425:686–91.

36. Breitkreutz A, Choi H, Sharom JR, Boucher L, Neduva V, Larsen B, et al. A global protein kinase and phosphatase interaction network in yeast. Science. 2010;328:1043–6.

37. Sunter J, Wickstead B, Gull K, Carrington M. A new generation of T7 RNA polymerase-independent inducible expression plasmids for Trypanosoma brucei. PLoS ONE. 2012;7:e35167.

38. Hammond MJ, Nenarokova A, Butenko A, Zoltner M, Dobáková EL, Field MC, et al. A uniquely complex mitochondrial proteome from Euglena gracilis. Ãvila-Arcos MC, editor. Mol. Biol. Evol. 2020;66:4.

39. Ogris E, Mudrak I, Mak E, Gibson D, Pallas DC. Catalytically inactive protein phosphatase 2A can bind to polyomavirus middle tumor antigen and support complex formation with pp60(c-src). J Virol. 1999;73:7390–8.

40. Nübel G, Sorgenfrei FA, Jäschke A. Boronate affinity electrophoresis for the purification and analysis of cofactor-modified RNAs. Methods. 2017;117:14–20.

41. Igloi GL, Kössel H. Affinity electrophoresis for monitoring terminal phosphorylation and the presence of queuosine in RNA. Application of polyacrylamide containing a covalently bound boronic acid. Nucleic Acids Res. 1985;13:6881–98.

42. Furuichi Y. Discovery of m(7)G-cap in eukaryotic mRNAs. Proc Jpn Acad Ser B Phys Biol Sci. 2015;91:394–409.

43. Kiledjian M. Eukaryotic RNA 5’-End NAD+ Capping and DeNADding. Trends Cell Biol. 2018;28:454–64.

44. Wang J, Alvin Chew BL, Lai Y, Dong H, Xu L, Balamkundu S, et al. Quantifying the RNA cap epitranscriptome reveals novel caps in cellular and viral RNA. Nucleic Acids Res. 2019;47:e130.

45. Luciano DJ, Vasilyev N, Richards J, Serganov A, Belasco JG. A Novel RNA Phosphorylation State Enables 5′ End-Dependent Degradation in Escherichia coli. Mol Cell. Elsevier Inc; 2017;67:44–6.

46. Chen YG, Kowtoniuk WE, Agarwal I, Shen Y, Liu DR. LC/MS analysis of cellular RNA reveals NAD-linked RNA. Nat Chem Biol. Nature Publishing Group; 2009;5:879–81.

47. Kowtoniuk WE, Shen Y, Heemstra JM, Agarwal I, Liu DR. A chemical screen for biological small molecule-RNA conjugates reveals CoA-linked RNA. Proc Natl Acad Sci USA. National Academy of Sciences; 2009;106:7768–73.

48. Clouet-d’Orval B, Batista M, Bouvier M, Quentin Y, Fichant G, Marchfelder A, et al. Insights into RNA processing pathways and associated-RNA degrading enzymes in Archaea. FEMS Microbiol Rev. 2018.

49. Kramer S, McLennan AG. The complex enzymology of mRNA decapping: Enzymes of four classes cleave pyrophosphate bonds. WIREs RNA. 2019;10:e1511.

50. Dunckley T, Parker R. The DCP2 protein is required for mRNA decapping in Saccharomyces cerevisiae and contains a functional MutT motif. EMBO J. 1999;18:5411–22.

51. Sharma S, Grudzien-Nogalska E, Hamilton K, Jiao X, Yang J, Tong L, et al. Mammalian Nudix proteins cleave nucleotide metabolite caps on RNAs. Nucleic Acids Res. 2020;48:6788–98.

52. Grudzien-Nogalska E, Kiledjian M. New insights into decapping enzymes and selective mRNA decay. WIREs RNA. Wiley-Blackwell; 2017;8:e1379.

53. Grudzien-Nogalska E, Wu Y, Jiao X, Cui H, Mateyak MK, Hart RP, et al. Structural and mechanistic basis of mammalian Nudt12 RNA deNADding. Nat Chem Biol. 2019;15:575–82.

54. Deana A, Celesnik H, Belasco JG. The bacterial enzyme RppH triggers messenger RNA degradation by 5’ pyrophosphate removal. Nature. 2008;451:355–8.

55. Cahová H, Winz M-L, Höfer K, Nübel G, Jäschke A. NAD captureSeq indicates NAD as a bacterial cap for a subset of regulatory RNAs. Nature. Nature Publishing Group; 2015;519:374–7.

56. Sasaki M, Takegawa K, Kimura Y. Enzymatic characteristics of an ApaH-like phosphatase, PrpA, and a diadenosine tetraphosphate hydrolase, ApaH, from Myxococcus xanthus. FEBS Lett. Federation of European Biochemical Societies; 2014;588:3395–402.

57. McLennan AG. Substrate ambiguity among the nudix hydrolases: biologically significant, evolutionary remnant, or both? Cell Mol Life Sci. 2012;70:373–85.

58. Bessman MJ, Walsh JD, Dunn CA, Swaminathan J, Weldon JE, Shen J. The gene ygdP, associated with the invasiveness of Escherichia coli K1, designates a Nudix hydrolase, Orf176, active on adenosine (5“)-pentaphospho-(5”)-adenosine (Ap5A). J Biol Chem. American Society for Biochemistry and Molecular Biology; 2001;276:37834–8.

59. Abdelraheim SR, Spiller DG, McLennan AG. Mammalian NADH diphosphatases of the Nudix family: cloning and characterization of the human peroxisomal NUDT12 protein. Biochem J. 2003;374:329–35.

60. Abdelraheim SR, Spiller DG, McLennan AG. Mouse Nudt13 is a Mitochondrial Nudix Hydrolase with NAD(P)H Pyrophosphohydrolase Activity. Protein J. Springer US; 2017;36:425–32.

61. van Dijk E, Cougot N, Meyer S, Babajko S, Wahle E, Séraphin B. Human Dcp2: a catalytically active mRNA decapping enzyme located in specific cytoplasmic structures. EMBO J. 2002;21:6915–24.

62. Xu J, Yang JY, Niu QW, Chua NH. Arabidopsis DCP2, DCP1, and VARICOSE Form a Decapping Complex Required for Postembryonic Development. Plant Cell. 2006;18:3386–98.

63. Williams CW, Elmendorf HG. Identification and analysis of the RNA degrading complexes and machinery of Giardia lamblia using an in silico approach. BMC Genomics. BioMed Central; 2011;12:586–15.

64. Kumar S, Stecher G, Tamura K. MEGA7: Molecular Evolutionary Genetics Analysis Version 7.0 for Bigger Datasets. Mol. Biol. Evol. 2016;33:1870–4.

65. Brun R, Schönenberger. Cultivation and in vitro cloning or procyclic culture forms of Trypanosoma brucei in a semi-defined medium. Short communication. Acta Trop. 1979;36:289–92.

66. McCulloch R, Vassella E, Burton P, Boshart M, Barry JD. Transformation of monomorphic and pleomorphic Trypanosoma brucei. Methods Mol Biol. 2004;262:53–86.

