## Supplementary Figure S1 and S2 for "Is mRNA decapping activity of ApaH like phosphatases (ALPH’s) the reason for the loss of cytoplasmic ALPH’s in all eukaryotes but Kinetoplastida?"

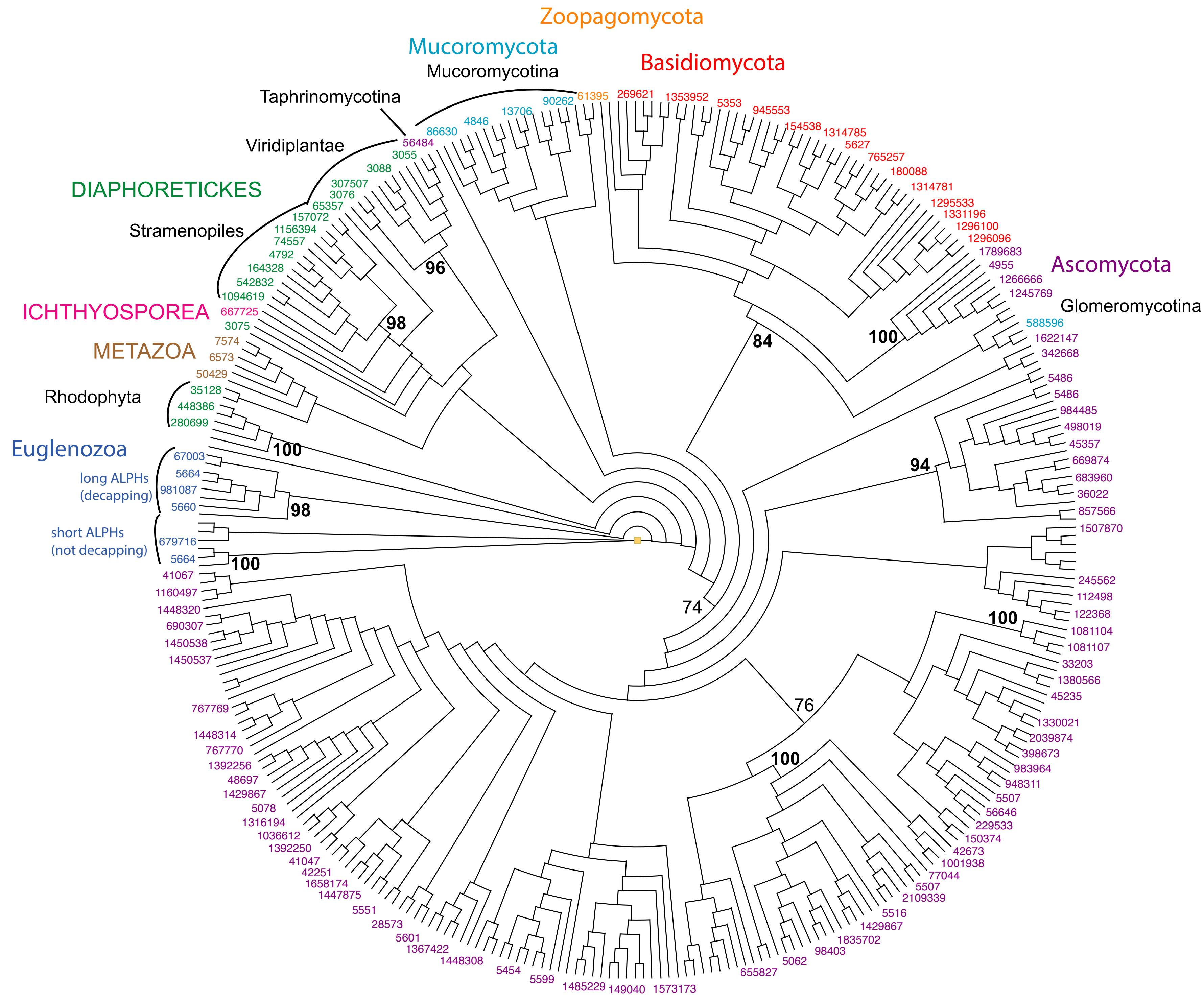


**Figure S1: Maximum Likelihood tree of ALPH in eukaryotes.**
This tree was built from catalytic domains (insertions removed) of 384 ALPH proteins; all sequences are liststed in Supplementary Table S1b.The PhyML tree was inferred using substitution models determined by Prot-Test (1). Boot- strapping was performed for 1000 replicates and the best tree topology was inferred. Bootstrap values above 80 are significant.


(1) Abascal, F., Zardoya, R. & Posada, D. ProtTest: selection of best-fit models of protein evolution. Bioinformatics 21, 2104–2105 (2005).

Figure S2

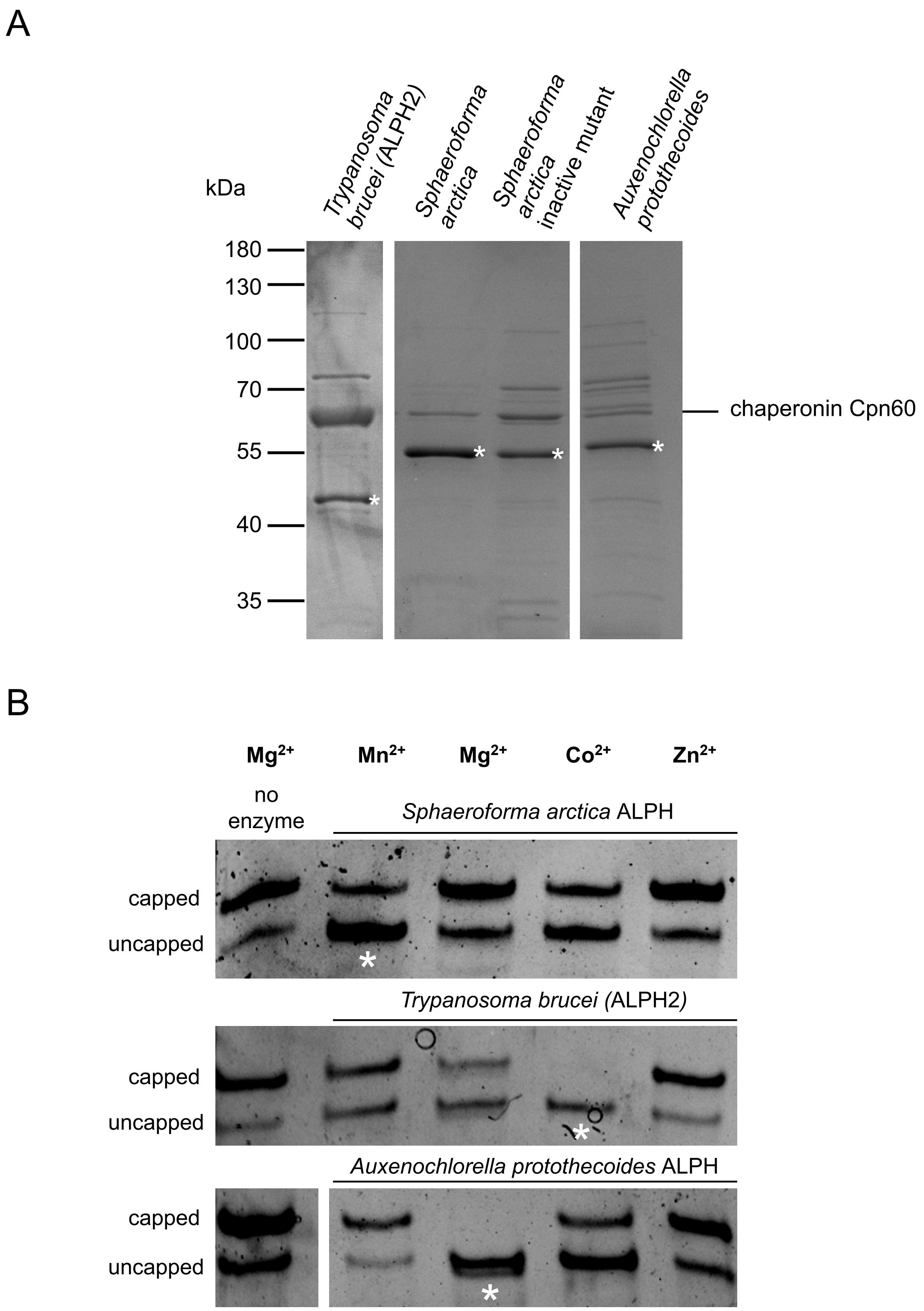


**Figure S2: Recombinant ALPH proteins: production and ion requirements**

**(A)** Recombinant ALPH proteins were produced in Arctic Express DE cells and purified via Nickel affinity chromatography. A Coomassie stained gel loaded with the purified proteins (marked by askterisks) is shown. Note that the Arctic Express DE cells contain chaperons. Chaperonin Cpn60 is often co-purified, in the case of TbALPH2 to a large extent.

**(B)** The enzymes were tested for *in vitro* decapping activity in the presence of 1 mM of divalent ions, as indicated. A reaction without enzyme served as negative control. For each enzyme, an asterisk marks the conditions with best decapping activity. The gels are representative gels out of at least three replicates.
